# Ultrasound-Mediated Gene Therapy in Alzheimer’s Disease Validated through In Vivo PET Imaging

**DOI:** 10.64898/2026.02.02.703398

**Authors:** Yutong Guo, Josquin Foiret, Javier Ajenjo, Rim Malek, Nisi Zhang, Jai Woong Seo, James Wang, Basit Latief Jan, Marina Nura Raie, Spencer K. Tumbale, Corinne Beinat, Katherine W. Ferrara

## Abstract

Efficient, spatially selective delivery of adeno-associated virus (AAV) therapeutics to deep brain structures remains a major challenge to gene therapy for Alzheimer’s disease (AD), owing to limited transport across the blood-brain barrier (BBB) and poor penetration to target neurons. Here, we establish an integrated, noninvasive imaging and therapy platform that combines microbubble-enhanced focused ultrasound (MB-FUS) with positron emission tomography/computed tomography (PET/CT) to transiently modulate the BBB, enhance region-specific AAV delivery following systemic dosing, and longitudinally track transduction in vivo. Optimized MB-FUS achieved targeted hippocampal delivery of systemically administered AAV9 in healthy mice, resulting in a 10-fold enhancement of neuronal transduction as compared to non-FUS controls. Importantly, longitudinal PET reporter gene imaging in the 5xFAD AD model demonstrated robust brain AAV transduction that remained stable for at least seven months. Finally, to assess therapeutic impact, we used brain-derived neurotrophic factor (BDNF) as a test cargo. MB-FUS-facilitated delivery elevated BDNF expression in targeted regions and produced short-term improvements in synaptic signaling in 5xFAD mice. Collectively, these results highlight MB-FUS as a next-generation delivery platform to overcome barriers to AAV therapeutic delivery in Alzheimer’s disease and position longitudinal PET assessment as a critical, translatable tool for monitoring and optimizing gene therapy.

## Introduction

Alzheimer’s disease (AD), the most common form of dementia, remains a major unmet medical challenge, affecting over 32 million people worldwide.^1–3^ Although recent FDA approvals of amyloid-targeting drugs (e.g., aducanumab, lecanemab, donanemab) have sparked hope, their modest cognitive benefit and potential side effects underscore the need for approaches that act beyond amyloid.^3–5^ Gene therapy offers a complementary, mechanism-diverse strategy by delivering transgenes that promote neuroprotection, restore synaptic and circuit function, enhance autophagy-mediated clearance of disease-related proteins, and modulate disease-associated pathways.^6–8^ Among available platforms, adeno-associated virus (AAV) vectors are particularly attractive for central nervous system (CNS) applications owing to their high transduction efficiency, low immunogenicity, and capacity for long-term transgene expression.^7,9,10^

A major obstacle to systemic AAV delivery is the blood-brain barrier (BBB), a tightly regulated neurovascular interface that restricts macromolecular transport from the circulation into the CNS parenchyma.^11,12^ Several alternative delivery routes have been evaluated clinically but each fall short: stereotactic injection is invasive and limited by targeting error and restricted coverage within the intended structure, whereas intra cerebrospinal fluid (CSF) delivery offers limited spatial control and inefficient access to deep parenchyma.^11,12^ Engineered BBB-crossing capsids can increase CNS delivery in select settings but again lack spatial targeting.^10,13–17^ Beyond baseline BBB permeability, AD-associated pathology, including BBB dysfunction, reduced transporter expression (e.g., LRP1), amyloid accumulation, and neuroinflammation, introduces structural and functional heterogeneity that adds complexity to vector delivery.^18,19^ Whether this pathological heterogeneity facilitates or impedes AAV transduction remains unclear, as regional permeability changes may coexist with perivascular debris accumulation that restricts parenchymal access and functional effectiveness.^18^ Together, these constraints motivate a delivery strategy that is noninvasive and scalable while enabling efficient, region-specific vector delivery, robust neuronal transduction, and minimal off-target biodistribution and peripheral toxicity.

Microbubble-enhanced focused ultrasound (MB-FUS) offers a unique noninvasive approach for targeted drug delivery to the brain. Focused ultrasound (FUS) beams can be precisely directed through the intact skull to specific brain regions with sub-millimeter accuracy. When combined with intravenously administered microbubbles (MB; lipid- or albumin-shelled gas pockets, 1-10 μm diameter), low-intensity FUS induces controlled oscillations that exert mechanical stress in brain vessels, transiently increasing BBB permeability (4-24 hours)^20–26^ and facilitating interstitial transport^27,28^. This spatiotemporal control creates a therapeutic window for region-selective delivery, and several preclinical studies have demonstrated enhanced AAV transduction in targeted brain regions using MB-FUS.^25,26,29–39^ More importantly, completed and ongoing clinical trials have demonstrated that FUS-mediated BBB opening is safe and feasible for AD treatment^22–24,40–49^, and a recent study combining MRI-guided FUS with aducanumab reported greater amyloid-β reduction in targeted regions^50^, underscoring the platform’s translational potential for targeted gene therapy in AD.

Despite promising preclinical and clinical studies exploring MB-FUS-mediated AAV delivery, critical gaps limit translation and optimization. Variations across studies in AAV serotypes, dosages, FUS parameters, and target regions, without standardized quantification methods, make direct comparisons and protocol optimization challenging. Moreover, existing readouts for assessing AAV delivery have significant limitations. Endpoint histology requires tissue sacrifice and cannot provide whole-organ quantification, while gadolinium-enhanced MRI and CSF sampling provide only indirect evidence of BBB opening, not direct measures of transduction or sustained transgene expression. Positron emission tomography (PET) reporter gene imaging addresses this gap by offering whole-body, quantitative, and clinically translatable visualization of transgene expression with high sensitivity and temporal resolution.^51–53^ We previously demonstrated using PET imaging of radiolabeled AAV9 and PET reporter gene imaging that MB-FUS substantially enhances vector delivery and transgene expression in targeted brain regions of healthy mice.^51^ However, whether these findings translate to AD, where BBB dysfunction and pathological heterogeneity may unpredictably alter delivery efficiency and transduction, and whether improved delivery yields meaningful therapeutic efficacy remain unclear. Together, these limitations underscore the need for noninvasive, quantitative imaging tools that can longitudinally monitor transgene expression and assess transduction efficiency in diseased brains to guide protocol optimization and therapeutic translation.

Here, we establish an integrated noninvasive platform combining MB-FUS with PET reporter gene imaging to enable region-specific AAV delivery and longitudinal monitoring of transgene expression in an AD murine model. We first optimized MB-FUS parameters using mathematical modeling of microbubble dynamics and in vivo validation across two frequencies (0.8 MHz and 1.5 MHz) to maximize hippocampal targeting precision, delivery efficiency, and neuronal transduction, assessed via contrast-enhanced MRI, longitudinal fluorescent reporter imaging, and immunofluorescence. Critically, we then applied quantitative PET reporter gene imaging in both healthy and 5xFAD AD model mice to directly test whether parameters optimized in healthy brains maintain delivery and transduction efficiency in the context of AD-related BBB dysfunction and pathological heterogeneity, demonstrating robust and sustained transgene expression in targeted regions over seven months. Finally, to evaluate therapeutic potential, we delivered brain-derived neurotrophic factor (BDNF) via MB-FUS-enhanced AAV9 and assessed functional improvements in synaptic signaling and neuroprotection in the 5xFAD hippocampus. Together, this work validates MB-FUS as a scalable, clinically translatable strategy for targeted gene therapy in AD and positions quantitative PET imaging as a critical tool for optimizing delivery protocols across disease contexts and monitoring treatment efficacy in vivo.

## Results

### Clinically translatable multi-frequency FUS systems achieve region-specific and safe AAV transduction in the healthy brain

To optimize MB-FUS parameters for hippocampal targeting, we first confirmed that 5 µm monodisperse lipid-shelled microbubbles are more potent in increasing BBB permeability than 2 µm microbubbles, as assessed by contrast-enhanced MRI and dextran extravasation (Suppl. Method 1; Suppl. Fig. 1), consistent with prior reports.^54–59^ Therefore, we used 5 µm microbubbles throughout this work. Next, to identify a frequency that balances effective skull penetration with robust microbubble-mediated BBB modulation and is compatible with clinical transcranial FUS systems, we tested 0.8 MHz and 1.5 MHz in parallel (Suppl. Fig. 2). To mitigate differences in focal dimensions between frequencies, we used phased-array electronic steering to deliver volumetric sonication for 1.5 MHz (see Methods), enabling more direct frequency comparison. Our FUS system enables precise focal targeting with real-time monitoring of MB activity via passive acoustic mapping (PAM) during sonication (Suppl. Fig. 2). As expected, 0.8 MHz showed a reduced skull attenuation compared to 1.5 MHz (Fig. 1B), though insertion loss increased with mouse age, particularly in animals younger than 3 months, likely due to ongoing skull maturation (Fig. 1B). To predict how frequency influences 5 µm MB oscillation dynamics and resulting bioeffects within brain vasculature, we simulated MB oscillations within brain vasculature using a mathematical model based on the Marmottant model at two frequencies: 0.8 MHz and 1.5 MHz.^55,60^ The model predicted that MB excited at 0.8 MHz oscillate at relative greater amplitude compared to 1.5 MHz (ΔR/R_0_ of 16% vs. 3%, respectively) under the same acoustic pressures (Fig. 1C), suggesting that lower-frequency ultrasound may induce stronger mechanical effects on the vasculature and enhance therapeutic delivery. To test this prediction in vivo, we performed MB-FUS treatment targeting the right hippocampus, a brain region critical for memory and cognition and one of the earliest affected in AD. FUS was applied during intravenous microbubble injection (2.5 × 10^8^ MBs/kg) using a peak negative pressure of 320 kPa (Supple. Method 2; 1 ms pulses, 5 Hz PRF), calibrated for skull insertion loss at both frequencies. During sonication, we monitored MBs acoustic emissions using passive acoustic monitoring, confirming MB oscillations through the detection of signature harmonic signals (Fig. 1D; Suppl. Fig. 2). Moreover, we observed minimal broadband emissions at both frequencies throughout the treatment, indicating that MB underwent stable oscillations rather than inertial cavitation. This is critical for safety, as broadband emissions are associated with microbubble collapse and potential hemorrhaging (Fig. 1D). Ten minutes after MB-FUS treatment, T1-weighted magnetic resonance imaging (MRI) following intravenous injection of a gadolinium-based contrast agent confirmed increased vascular permeability in the hippocampus, as evidenced by contrast extravasation (Fig. 1E). This result validates both the enhanced permeability and the spatial precision of our targeting approach at both ultrasound frequencies setups. Moreover, histological analysis of brain slices showed that the BBB opening in the FUS-targeted areas did not result in hemorrhage, further demonstrating the safety of the procedure at this pressure (Fig. 1F).

**Fig. 1.**
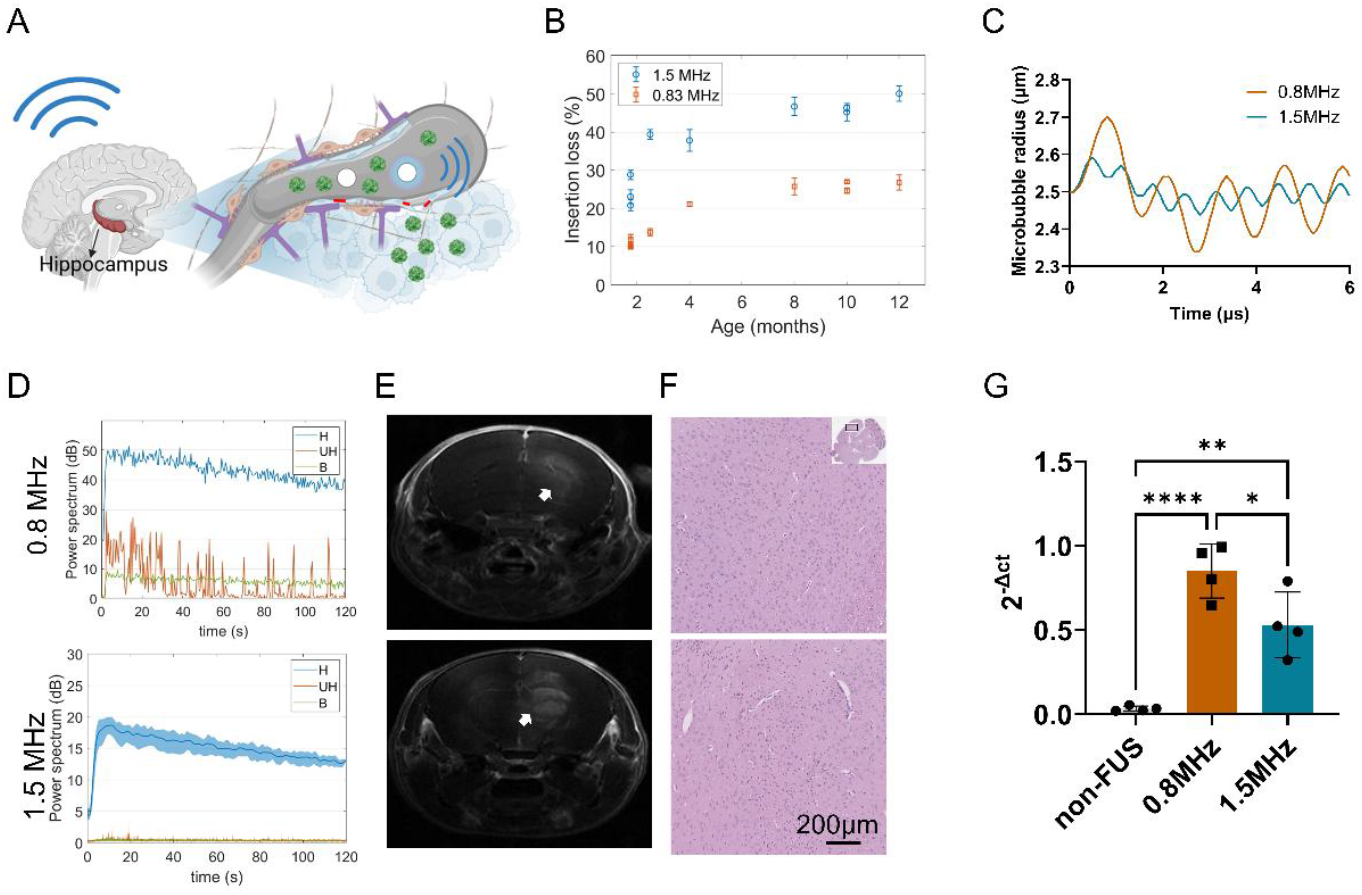
Multi-frequency ultrasound systems enable BBB opening for targeted AAV delivery. A) Schematic of microbubble-enhanced focused ultrasound for improved targeted AAV delivery to the hippocampus. B) Measurement of ultrasound insertion loss through the mouse skull at 0.8 MHz and 1.5 MHz from 2 to 12 months of age. C) Mathematical modeling of microbubble oscillations inside a 10 μm vessel excited with 0.8 MHz and 1.5 MHz frequencies at 175kPa peak negative pressure. D) Representative power levels of acoustic emissions originating from MB oscillations during a typical 120s treatment using 0.8 MHz (upper) and 1.5 MHz (lower) frequency. Emissions are processed in the frequency domain in specific bands (H: harmonics, UH: ultra-harmonics, B: broadband). E) Representative contrast enhanced T1-weighted MR image 10 minutes after MB-FUS treatment using 0.8 MHz (upper) and 1.5 MHz (lower) frequency. F) Representative Hematoxylin-eosin (H&E) staining images of the brain treated with 0.8 MHz (upper) and 1.5 MHz (lower) frequency. G) Reverse transcription quantitative polymerase chain reaction (RT-qPCR) assessment of ssDNA of AAV9 transgene (tdTomato) in brain hemispheres at 21 hours after the treatment. P-values were determined by one-way analysis of variance (ANOVA) and were adjusted using Bonferroni correction. Plots show mean ± S.D (n = 4). *P ≤ 0.05, **P ≤ 0.01, **** P ≤ 0.0001.

After confirming targeting precision, we assessed the ability of MB-FUS to enhance delivery of intravenously administered AAV9 (3 × 10^13^ vg/kg), selected for its established CNS tropism following systemic administration, to the brain using both frequencies. MB-FUS was started immediately following systemic AAV9 injection. Quantitative PCR analysis of AAV9 single-stranded DNA copies in the sonicated hemisphere 21 hours post-treatment revealed significant increases in vector genome delivery: 25-fold at 0.8 MHz (p < 0.0001) and 15-fold at 1.5 MHz (p = 0.0028) compared to AAV-only controls (Fig. 1G). The greater enhancement at 0.8 MHz is consistent with the 5 µm MB oscillation amplitude predicted by our mathematical model. Together, these findings establish our multi-frequency FUS platform, particularly at 0.8 MHz, as a safe and effective strategy for region-specific AAV delivery in the healthy brain.

### Neuron-specific and liver-detargeted AAVs transduction achieved by MB-FUS revealed by fluorescent reporter imaging

Having established that our MB-FUS systems achieve enhanced, targeted AAV9 delivery, we next assessed transduction efficiency in healthy brains following MB-FUS treatment using a fluorescent reporter gene, tdTomato (Fig. 2A). In vivo optical imaging revealed that AAV9 injection alone produced moderate tdTomato expression in the brain compared to non-AAV, non-FUS controls. However, MB-FUS treatment at both 0.8 MHz and 1.5 MHz substantially enhanced transgene expression relative to AAV alone, with elevated levels sustained for up to 6 months (Fig. 2B-C; Suppl. Fig. 3). Notably, while both frequencies yielded comparable transduction within the first month, the 0.8 MHz group demonstrated consistently higher tdTomato expression thereafter, indicating a potential long-term advantage (Fig. 2C; Suppl. Fig. 3). Ex vivo brain analysis at 1 and 7 months after treatment confirmed these findings, with significantly enhanced expression in both MB-FUS groups compared to non-FUS controls and higher transduction with 0.8 MHz at 7 months (Fig. 2D-F).

**Fig. 2.**
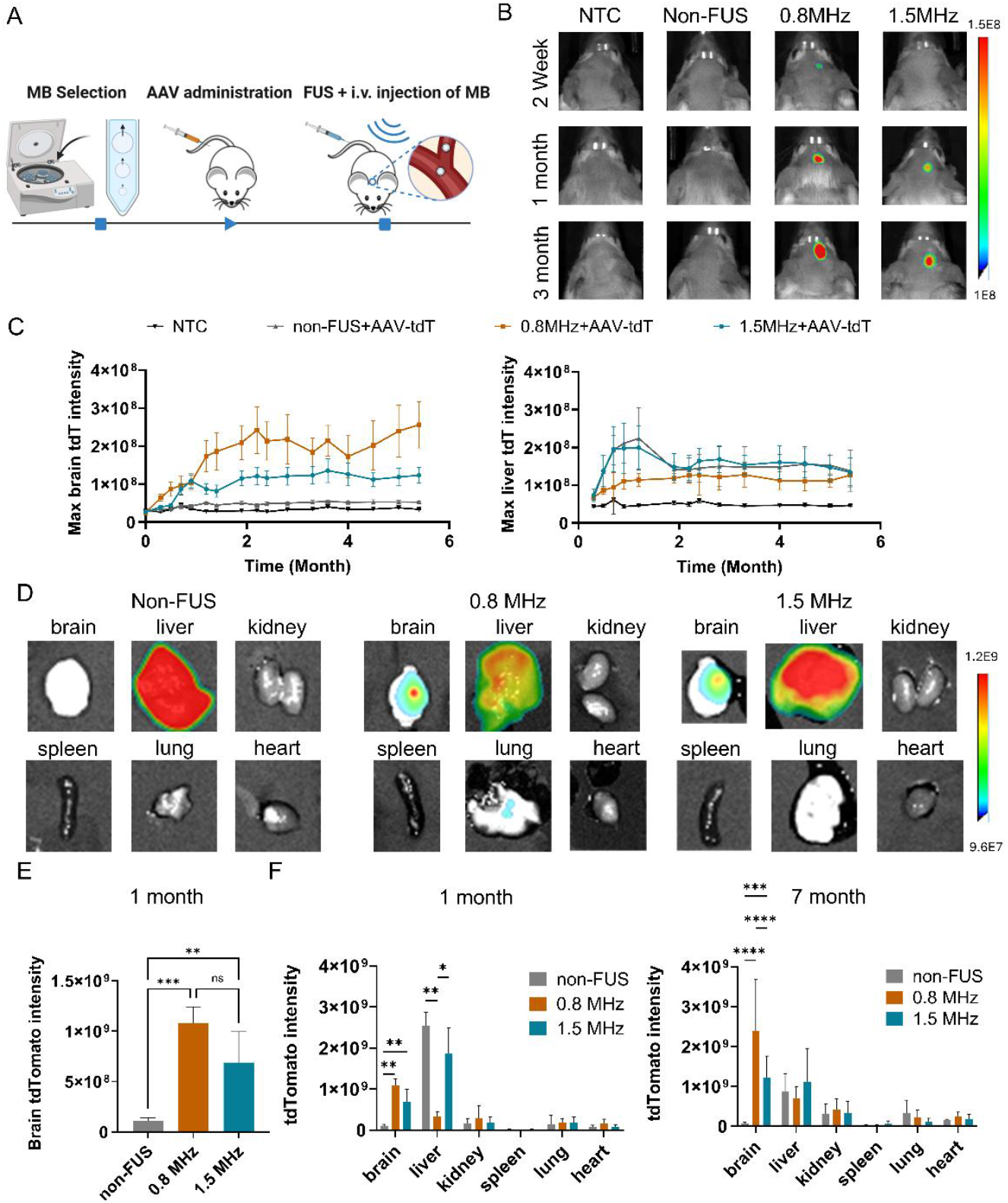
In vivo fluorescent reporter gene imaging reveals enhanced and stable brain-targeted AAV9 transduction with liver detargeting following MB-FUS treatment. A) In vivo experimental protocol for AAV delivery with MB-FUS. B) Representative in vivo imaging of AAV9-tdTomato expression in the brain at 2 weeks, 1 month, and 3 months after treatment. C) Quantification of in vivo imaging of maximum tdTomato expression in the brain (left) and liver (right) over a 6-month period following treatment. D) Representative ex vivo images of tdTomato transduction biodistribution 1 month after treatment. E) Quantification of ex vivo imaging of tdTomato expression in the brain 1 month after treatment. P-values were determined by one-way analysis of variance (ANOVA) and were adjusted using Bonferroni correction. F) Quantification of ex vivo imaging of tdTomato transduction biodistribution 1 month (left) and 7 months (right) after treatment. P-values were determined by two-way analysis of variance (ANOVA) and were adjusted using Bonferroni correction. Plots show mean ± S.D (n=5). *P ≤ 0.05, **P ≤ 0.01, ***P ≤ 0.001. Abbreviations: tdT: tdTomato.

Interestingly, MB-FUS treatment markedly reduced off-target AAV9 expression in the liver, potentially addressing a major challenge in systemic AAV gene therapy, where AAV liver tropism raises concerns about hepatotoxicity, immune responses, and dose-limiting toxicity (Fig. 2C; Suppl. Fig. 4).^61^ In contrast to the sustained transduction observed in the brain, liver expression in AAV9-treated groups peaked at 1 month and subsequently declined. Unexpectedly, both in vivo and ex vivo optical imaging revealed that liver tdTomato expression was selectively reduced in the 0.8 MHz MB-FUS group compared to other treatment groups, with the most pronounced reduction observed between 1 and 2 months post-treatment (Fig. 2C-D, F; Suppl. Fig. 4). Quantitative analysis at 1 month confirmed that while the liver was the primary site of AAV9 accumulation in non-FUS controls (liver-to-brain ratio: 25 ± 4.6; maximum intensity), 0.8 MHz MB-FUS reversed this biodistribution, achieving a brain-to-liver ratio of 3.6 ± 0.8 (p = 0.013 vs. non-FUS; Fig. 2F). Although further investigation is needed to elucidate the underlying mechanism, these findings suggest that MB-FUS not only enables region-specific AAV9 delivery but may also reduce peripheral exposure and systemic toxicity by limiting off-target hepatic accumulation.

To evaluate the spatial and cell-type specificity of AAV9 transduction, we performed immunofluorescence analysis on brain sections harvested at 7 months post-treatment. Confocal microscopy revealed robust tdTomato expression in the hippocampus of MB-FUS-treated mice at both 0.8 MHz and 1.5 MHz, confirming effective transduction and precise regional targeting (Fig. 3A, C). Consistent with our prior observations^51^, we also detected transgene expression in the contralateral (non-sonicated) hippocampus, suggesting potential vector diffusion or axonal transport via interhemispheric connections following MB-FUS-mediated delivery^62^ (Fig. 3A). To elucidate cellular specificity, we quantified colocalization of AAV transduction (tdTomato) with neurons (NeuN) and vasculature (lectin), revealing a marked increase in neuronal transduction in MB-FUS-treated animals at both frequencies (0.8 MHz: 37.4% ± 6.5%; 1.5 MHz: 36% ± 9.1%) compared to the AAV-only group (4.2% ± 0.4%, p < 0.05), representing a 9-fold enhancement (Fig. 3B, D). Conversely, the majority of transduced cells in the AAV-only group were associated with blood vessels (28% ± 2.3% lectin^+^ vs. 4.2% ± 0.4% NeuN^+^; Fig. 3D-E), indicating limited parenchymal penetration and perivascular confinement. These findings demonstrate that MB-FUS overcomes vascular barriers to facilitate AAV9 extravasation into the brain parenchyma, thereby enhancing neuronal transduction efficiency. This shift from endothelial-restricted to neuron-preferential targeting underscores the capacity of MB-FUS to modulate vector biodistribution at the cellular level and improve therapeutic specificity in CNS gene delivery.

**Fig. 3.**
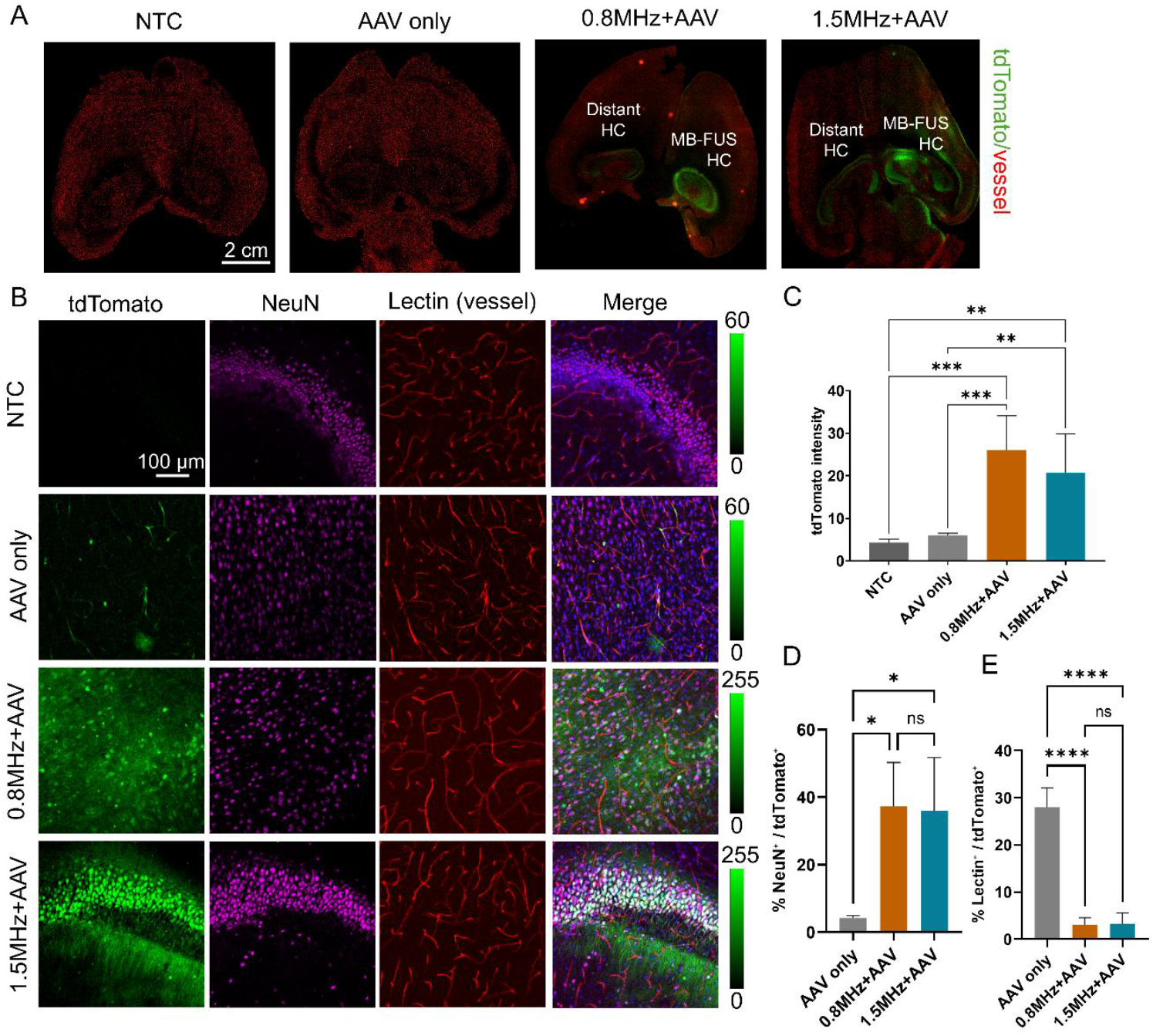
Immunofluorescent microscopy demonstrates that MB-FUS-enhanced delivery enables spatially targeted and neuron-specific AAV transduction in the brain. A) Representative fluorescent microscopy images showing AAV9-tdTomato transduction in the brains of healthy mice at 6 months post-treatment. From left to right: untreated control, MB-FUS only, AAV only, AAV + MB-FUS at 0.8 MHz, and AAV + MB-FUS at 1.5 MHz. B) Representative immunofluorescent staining of neurons and blood vessels at 6 months post-treatment, demonstrating cell-specific AAV transduction. Green: tdTomato; Magenta: NeuN (neuronal nuclei); Red: Lectin (vasculature); Blue: DAPI (nuclei). C) Quantification of brain tdTomato expression across treatment groups at 6 months post-treatment. (n = 4) D) Quantification of neuron-specific transduction, expressed as the percentage of NeuN^+^ cells among all tdTomato^+^ cells in the brain at 6 months post-treatment. (n = 3) E) Quantification of vessel cell-specific transduction, expressed as the percentage of lectin^+^ cells among all tdTomato^+^ cells in the brain at 6 months post-treatment. (n = 3) P-values were determined by one-way analysis of variance (ANOVA) and were adjusted using Bonferroni correction. Plots show mean ± S.D. n.s. no significance, P >0.05, *P ≤ 0.05, **P ≤ 0.01, ***P ≤ 0.001, **** P ≤ 0.0001. Abbreviations: HC: Hippocampus.

### PET/CT reporter gene imaging enables longitudinal in vivo monitoring of gene expression and reveals enhanced AAV9 transduction following MB-FUS treatment in an Alzheimer’s disease mouse model

While fluorescent reporter gene imaging provides valuable information in small animal models, its limited tissue penetration, sensitivity, and quantification accuracy restrict its translational potential. In contrast, PET/CT imaging offers deep tissue penetration, high sensitivity, and quantitative capability, making it a clinically translatable modality for evaluating AAV transduction efficiency and spatial distribution. Importantly, PET enables direct comparison of MB-FUS performance across healthy and diseased brains, addressing the critical question of whether AD-related BBB dysfunction and pathological heterogeneity alter the transduction efficiency. To leverage these advantages, we developed a PET-based method to assess MB-FUS-enhanced AAV9 transduction in both healthy and 5xFAD mice. Specifically, we engineered AAV9 vectors encoding the PET reporter gene pyruvate kinase M2 (PKM2) and employed [^18^F]DASA-10, a BBB-permeable radiotracer^51,53^, to noninvasively monitor and quantify PKM2 expression in vivo.

Following the same treatment protocol, mice received MB-FUS and AAV9-PKM2 delivery at three months of age (Fig 2A). Six months post-treatment, [^18^F]DASA-10 PET imaging revealed significantly enhanced AAV9-PKM2 expression in the sonicated brain regions of both healthy and 5xFAD mice (Fig 4; Suppl. Fig. 5). In healthy mice, standardized uptake values (SUVs) in the targeted regions were 0.64 ± 0.08 for the 0.8 MHz group and 0.63 ± 0.27 for the 1.5 MHz group, both significantly higher than the 0.20 ± 0.06 observed in non-FUS controls (p = 0.0003 and p = 0.0005, respectively) (Fig 4A-B). More importantly, 5xFAD mice showed similar trend of increased SUVs of 0.89 ± 0.13 (0.8 MHz) and 0.84 ± 0.12 (1.5 MHz), compared to 0.49 ± 0.07 in non-FUS controls (p = 0.0004 and p = 0.0028, respectively), indicating enhanced and sustained transgene expression in diseased brain environment with MB-FUS (Fig. 4D-E). Notably, SUVs were consistently higher in 5xFAD mice than in healthy mice across all groups, potentially reflecting both increased baseline permeability in the diseased vasculature and endogenous PKM2 upregulation associated with AD pathology^63^. qPCR analysis of PKM2 mRNA levels in the treated hemisphere confirmed the PET imaging findings. In healthy mice, MB-FUS showed higher expression with 25-fold (0.8 MHz) and 16-fold (1.5 MHz) increases vs. non-FUS controls (p < 0.003 for both frequencies; Fig. 4C). In 5xFAD mice, enhancement was even more pronounced with 0.8 MHz, yielding a 45-fold increase in expression (p = 0.0017; Fig. 4F). Across both mouse models, 0.8 MHz yielded significantly higher expression than 1.5 MHz (p < 0.05), consistent with the frequency-dependent enhancement observed in optical imaging quantification. Together, these results validate PET imaging as a noninvasive and clinically translatable approach for tracking AAV transduction and demonstrate effective, region-specific gene delivery with MB-FUS in both healthy and AD (5xFAD) mouse models.

**Fig. 4.**
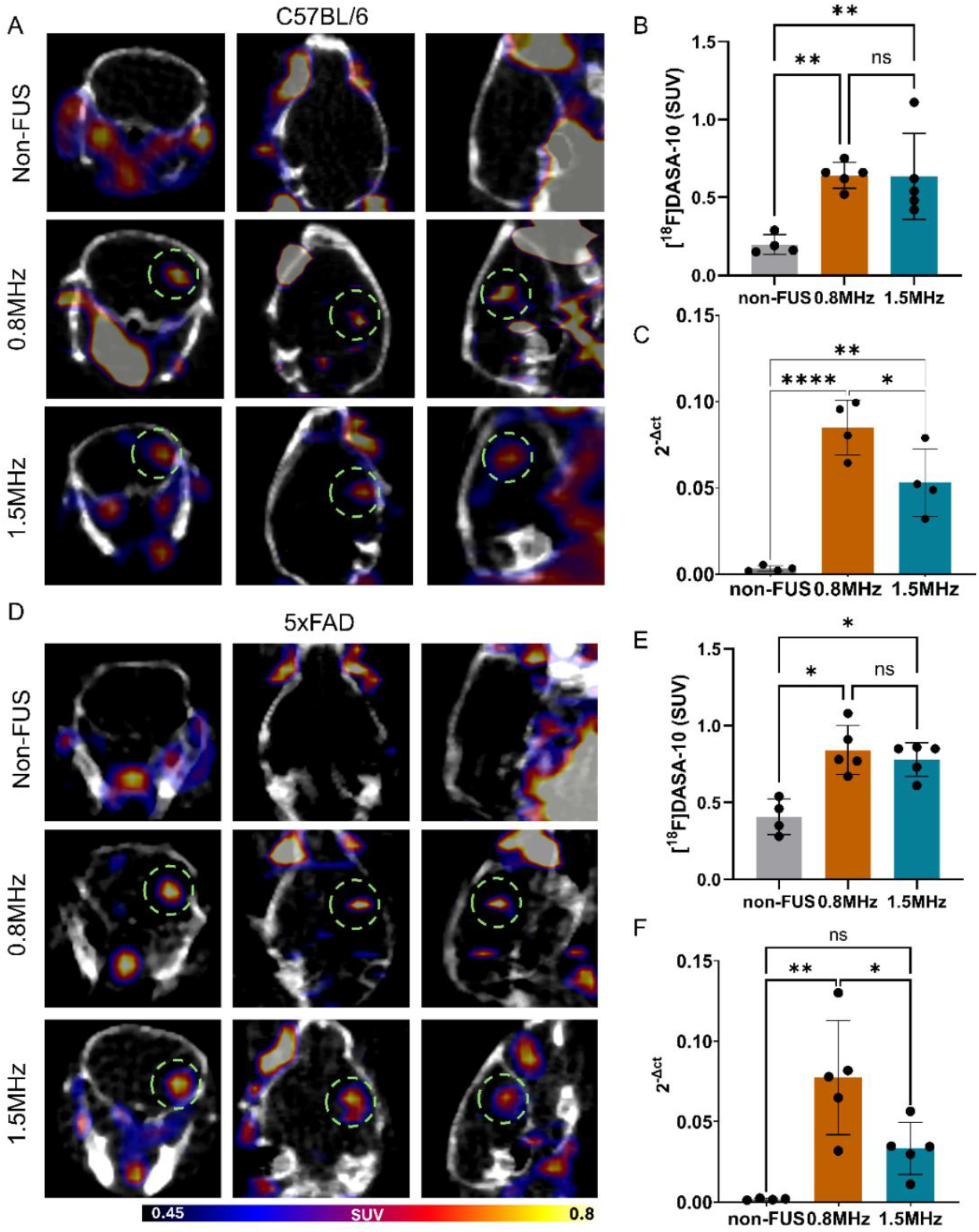
In vivo PET/CT imaging with [^18^F]DASA-10 quantifies AAV9-mediated PKM2 transduction following MB-FUS in both healthy and 5xFAD Alzheimer’s model mice. A) Representative PET/CT images (coronal, axial, sagittal) obtained immediately following [^18^F]DASA-10 administration, 6 months after MB-FUS-assisted AAV9-PKM2 delivery in healthy mice. Regions of MB-FUS treated region are marked with dotted green circles. B) Image-based quantification of standardized uptake values (SUVs) in regions of interest across treatment groups in healthy mice following [^18^F]DASA-10 injection. C) Reverse transcription quantitative polymerase chain reaction (RT-qPCR) quantification of PKM2 mRNA levels in MB-FUS-treated brain hemispheres of healthy mice 6 month following MB-FUS-mediated AAV9-PKM2 delivery. D) Representative PET/CT images (coronal, axial, sagittal) obtained immediately following [^18^F]DASA-10 administration, 6 months after MB-FUS-assisted AAV9-PKM2 delivery in 5xFAD mice. Regions of MB-FUS treated region are marked with dotted green circles. E) Image-based quantification of standardized uptake values (SUVs) in regions of interest across treatment groups in 5xFAD mice following [^18^F]DASA-10 injection. F) RT-qPCR quantification of PKM2 mRNA levels in MB-FUS-treated brain hemispheres of 5xFAD mice 6 month following MB-FUS-mediated AAV9-PKM2 delivery. P-values were determined by one-way analysis of variance (ANOVA) and were adjusted using Bonferroni correction. Plots show mean ± S.D. (n = 4-5) n.s. no significance, P >0.05, *P ≤ 0.05, **P ≤ 0.01, **** P ≤ 0.0001.

### MB-FUS-enhanced AAV9:BDNF delivery produces transient synaptic improvements in 5xFAD mice

To evaluate the therapeutic potential of MB-FUS-mediated gene delivery, we assessed whether enhanced hippocampal delivery of AAV9 encoding brain-derived neurotrophic factor (BDNF) could improve synaptic function in 5xFAD mice. BDNF was selected as a therapeutic payload due to its established capacity to reduce neuronal death, enhance synaptic signaling, and improve hippocampal-dependent memory in preclinical AD models^64–69^, which has led to its recent advancement into a first-in-human AAV-based gene therapy trial (NCT05040217). Quantitative PCR confirmed robust BDNF transgene expression following MB-FUS treatment, with 50-112-fold improvement vs. non-FUS controls observed at 1 and 7 months post-treatment (p < 0.05; Fig. 5A, E). Importantly, at 1-month post-treatment, MB-FUS-treated animals showed a significant increase in the presynaptic marker synapsin 1 (Syn1) compared to both AAV-only and non-treatment controls (Fig. 5B), alongside trends toward elevated postsynaptic markers synaptophysin (Syp) and PSD-95 (DLG4), though not statistically significant (Fig. 5C–D). By 7 months, synaptic marker levels had returned to baseline across all treatment groups (Fig. 5F–H). Together, these findings provide proof-of-concept that MB-FUS-enhanced AAV9-BDNF delivery can elicit early synaptic improvements in 5xFAD mice, and suggest that optimized dosing, earlier intervention, or sustained expression strategies may be required to achieve long-term functional rescue in AD models.

**Fig. 5.**
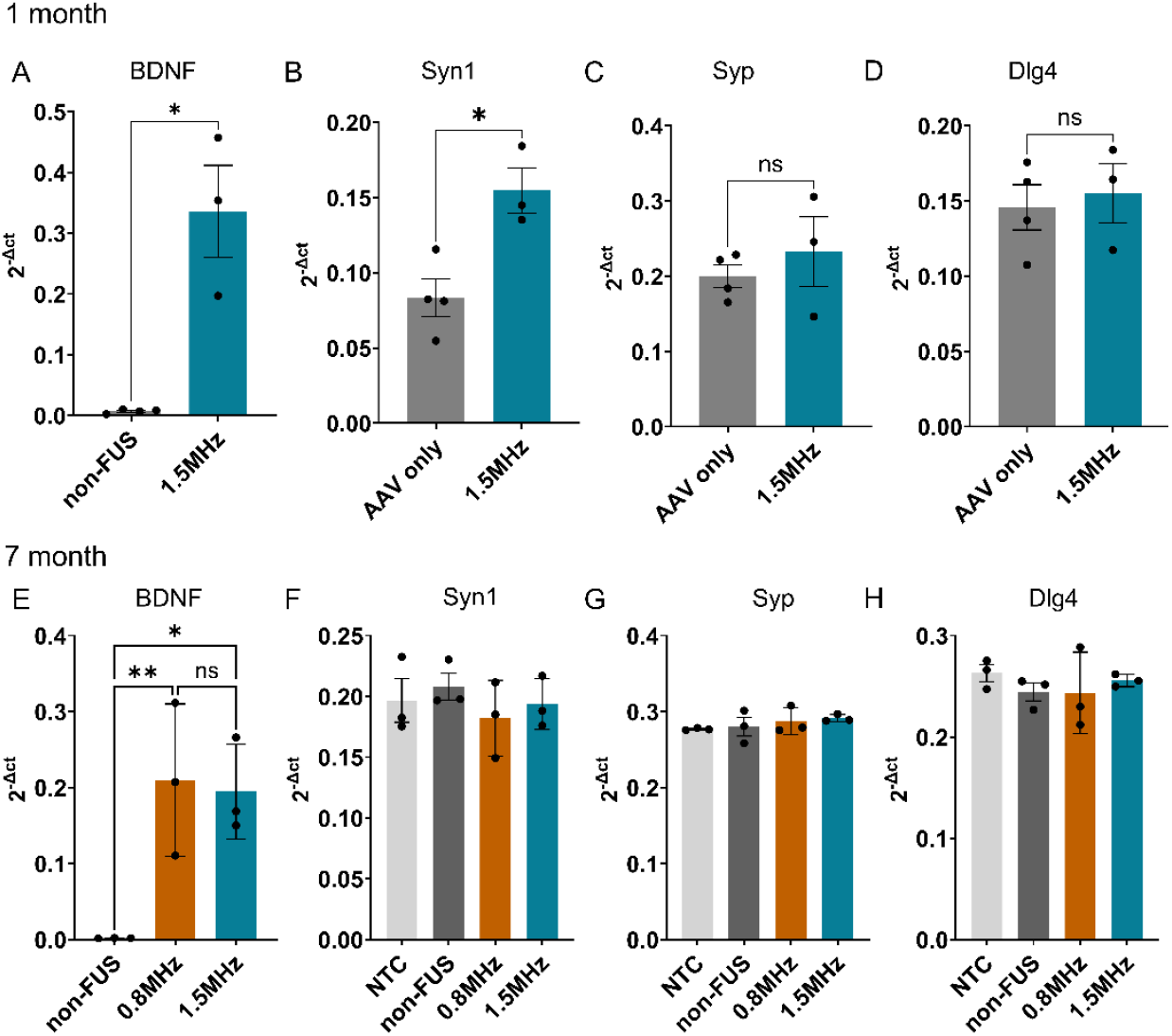
MB-FUS-enhanced AAV9-BDNF delivery produces transient synaptic improvements in 5xFAD mice. A) Reverse transcription quantitative polymerase chain reaction (RT-qPCR) quantification of BDNF mRNA levels in the treated hemisphere of 5xFAD mice 1 month following MB-FUS-mediated AAV9-BDNF delivery. RT-qPCR quantification of the B) presynaptic marker synapsin 1 (Syn1), C-D) postsynaptic markers synaptophysin (Syp) and PSD-95 (Dlg4) one month post treatment. E) RT-qPCR quantification of BDNF mRNA levels in the hippocampus 7 months following MB-FUS-mediated AAV9-BDNF delivery. RT-qPCR quantification of synaptic markers F) Syn1, G) Syp, and H) Dlg4 at 7 months post-treatment. P-values were determined by one-way analysis of variance (ANOVA) and were adjusted using Bonferroni correction. Plots show mean ± S.D. (n = 3) n.s. no significance P >0.05, *P ≤ 0.05, **P ≤ 0.01.

## Discussion

In this study, we established an integrated platform combining microbubble-enhanced focused ultrasound (MB-FUS) with positron emission tomography (PET) reporter gene imaging to enable noninvasive, region- and cell-specific AAV delivery and longitudinal monitoring of transgene expression in AD. Through systematic optimization, we identified 0.8 MHz as the preferred frequency for hippocampal targeting, achieving up to 25-fold enhancement in vector delivery, 9-fold improvement in neuronal transduction, and sustained expression over 7 months in healthy mice, with reduced off-target liver accumulation. Critically, quantitative PET imaging confirmed that delivery parameters optimized in healthy brains translated effectively to the AD context, demonstrating durable transgene expression despite BBB dysfunction and pathological heterogeneity. Moreover, proof-of-concept therapeutic testing with BDNF revealed early synaptic improvements, validating the platform’s capacity to deliver functional therapeutics. Together, these findings establish MB-FUS with PET monitoring as a clinically translatable strategy for targeted, quantifiable gene therapy in neurodegenerative disease.

A central finding of this work is the optimization of MB-FUS parameters to 0.8 MHz based on mathematical modeling and in vivo validation. Modeling predicted that 5 µm microbubbles driven at 0.8 MHz would oscillate at greater amplitude than at 1.5 MHz under equivalent acoustic pressures, which we confirmed experimentally through enhanced vector delivery, sustained neuronal transduction, and reduced hepatic off-target expression. Importantly, this frequency bridges a translational gap between preclinical and clinical practice. While preclinical studies typically employ frequencies above 1 MHz for smaller focal zones and better precision, clinical transcranial FUS systems operate at 200 kHz-1 MHz to minimize skull attenuation, enhance penetration, and reduce heating risk.^70^ Since FUS parameters critically influence bioeffects in a nonlinear manner^55^, using a frequency compatible with both preclinical models and clinical systems is essential for scalable translation. Our finding that 0.8 MHz achieves robust, neuron-specific hippocampal delivery while balancing penetration and spatial precision suggests that this frequency can bridge preclinical and clinical applications, allowing optimized protocols to translate directly to patients.

The enhanced delivery achieved by MB-FUS translated not only to increased overall transgene expression but also to a shift in cellular tropism, redirecting systemically administered AAV9 from predominantly perivascular and glial cells to robust neuronal transduction in adult mice. Our AAV-only controls showed less than 5% neuronal labeling, consistent with prior reports and providing further evidence that this limitation stems from perivascular trapping by mature astrocytic endfeet that restricts parenchymal penetration.^71–73^ Critically, this suggests that limited neuronal transduction reflects a delivery barrier that restricts extravasation and diffusion, rather than inherent capsid tropism. By transiently modulating BBB permeability and enhancing interstitial transport, MB-FUS overcomes perivascular entrapment, enabling deep neuronal access and converting a systemically scalable vector into a regionally targeted, neuron-specific platform.

A critical challenge for clinical translation is determining whether parameters optimized in healthy brains maintain efficacy in the pathological context of AD. To address this, we integrated quantitative PET reporter gene imaging, which provides noninvasive, longitudinal assessment of transgene expression and addresses a critical gap in the field by providing standardized methods to monitor AAV transduction in vivo. Prior preclinical studies have relied on endpoint histology or indirect readouts such as gadolinium-enhanced MRI, which provide limited quantitative information and cannot assess longitudinal transgene expression or compare delivery efficiency across cohorts and disease contexts. By employing PET imaging of the reporter gene PKM2 with the BBB-permeable radiotracer [^18^F]DASA-10, we demonstrated for the first time that MB-FUS parameters optimized in healthy mice translate effectively to the 5xFAD AD model, yielding robust and stable transgene expression over 7 months despite BBB dysfunction and pathological heterogeneity. Critically, molecular confirmation via qPCR revealed 45-112-fold enhancement in AAV transduction in the 5xFAD brain across reporter and therapeutic transgenes compared to non-FUS controls, suggesting that combining focal BBB modulation with systemic administration could enable substantially lower vector doses while maintaining therapeutic efficacy, thereby reducing systemic exposure, off-target toxicity, and manufacturing costs. Beyond technical validation, PET imaging offers translational potential by enabling systematic dose optimization, facilitating cross-laboratory protocol comparison, and providing a clinically compatible platform for real-time monitoring and individualized therapy adjustment in patients. Importantly, this approach is broadly applicable beyond Alzheimer’s disease to other CNS disorders requiring region-specific gene delivery, including Parkinson’s disease, epilepsy, and glioblastoma, positioning PET-guided MB-FUS as a generalizable strategy for precision gene therapy in the brain.

To assess therapeutic potential, we delivered AAV9-BDNF via MB-FUS in 5xFAD mice and observed sustained transgene expression over 7 months with early synaptic improvements, but these effects were not maintained at later timepoints. While BDNF has shown neuroprotective efficacy in prior preclinical studies^64–69^, our findings in the 5xFAD model revealed only transient benefit, highlighting that successful delivery does not automatically ensure sustained therapeutic efficacy. These results provide proof-of-concept that the MB-FUS platform can deliver functional therapeutics and elicit measurable biological responses in the AD brain, while also revealing key areas for optimization and further investigation. Several factors may contribute to the transient therapeutic effects observed. BDNF dose was not optimized in this study, and the relationship between vector dose, sustained BDNF expression, and long-term functional outcomes in the 5xFAD brain remains to be defined. Importantly, the PET reporter imaging platform established here enables noninvasive, longitudinal quantification of AAV transduction across different vector doses, facilitating systematic dose-titration studies to identify optimal dosing regimens that maximize therapeutic benefit, with direct translatability to individualized patients dosing in clinical trials. Moreover, the 5xFAD model is an aggressive, amyloid-driven system in which ongoing synapse loss may outpace BDNF-mediated rescue, particularly at later disease stages. Testing in tau-based, neuroinflammation-driven, or apolipoprotein E4 (APOE4) knock-in models will be essential to assess platform performance across the heterogeneity of AD pathology. Additional limitations include the mechanisms underlying the observed reduction in liver transduction at 0.8 MHz, which remain unclear and warrant further investigation. Potential explanations include altered AAV pharmacokinetics, increased brain accumulation reducing hepatic bioavailability, or frequency-dependent effects on systemic circulation, but rigorous biodistribution studies will be needed to distinguish these possibilities.

Collectively, our findings demonstrate that MB-FUS with PET monitoring addresses critical barriers to CNS gene therapy by enabling noninvasive, region-specific delivery with real-time quantification, and provides a scalable, clinically viable platform for therapeutic development in Alzheimer’s disease and beyond.

## Methods

### In vivo experiments

All animal experiments were conducted with a protocol approved by the Institutional Animal Care & Use Committee (IACUC) at Stanford University. Female mice aged 3 to 9 months were used in this study, with treatments initiated at 3 months of age and longitudinal monitoring until study endpoints at 9 months. C57BL/6 mice (Charles River Laboratories) were used to characterize BBB opening induced by multi-frequency MB-FUS. B6 albino mice (Jackson Laboratory) were used to assess AAV9-tdTomato transduction via in vivo and ex vivo optical imaging. To investigate AAV9 transduction and therapeutic efficacy in the context of AD, 5xFAD transgenic mice and age-matched B6SJLF1/J wild-type controls (Jackson Laboratory) were used. These cohorts were subjected to AAV9-mediated delivery of either PKM2 for PET/CT imaging or BDNF for functional therapeutic evaluation, with or without MB-FUS.

### Focused ultrasound system and transducer calibration

We developed custom multi-frequency FUS systems operating at 0.8 MHz and 1.5 MHz to investigate the effect of ultrasound frequency on MB-FUS-induced BBB opening and to assess whether a lower-frequency, which is more amenable to clinical translation, can support efficient and robust AAV-mediated gene delivery in the context of AD. The 0.8 MHz system consists of a single-element therapeutic transducer (H-115, Sonic Concepts, WA; fundamental frequency: 250 kHz; third harmonic frequency with matching network: 829 kHz) driven at 829 kHz with an arbitrary waveform generator (model 33500B, Agilent) and a 40-dB power amplifier (UltraX10, E&I) and a passive cavitation detector (5 MHz single element, unfocused, 3.175-mm aperture) oriented toward the acoustic focus of the therapeutic transducer with a 45° angle to reduce sensitivity to direct transmit beam reflection. The 1.5 MHz system, as previously described^51^, features a custom 128-element 2D therapeutic array (center frequency: 1.5 MHz) integrated with an L12-5 linear imaging transducer (38 mm aperture, Philips/ATL) positioned within the array’s central opening. A Vantage 256 ultrasound system (Verasonics) was used to do both image guidance, FUS therapy and acoustic emission recording during treatment. To account for smaller focal dimensions at 1.5 MHz, we expanded the treated region by raster scanning the 1.5 MHz focus over a 5 × 5 grid with 0.5 mm spacing (total sonicated volume ∼2.5 × 2.5 × 2.7 mm^3^ ≈ 17 mm^3^), allowing easier comparison between frequencies. Real-time analysis of acoustic emissions via passive cavitation detection (PCD) was employed to characterize MB oscillations throughout sonication for both frequencies. With the 1.5 MHz setup, the imaging transducer was also used to generate passive acoustic maps to localize acoustic emissions.^74^ This feedback mechanism allowed us to monitor cavitation dynamics and ensure sonications remained within predefined safety thresholds. The mechanical index (MI) during imaging was maintained at or below 0.35 to ensure safety.

The therapeutic transducers were calibrated both in free-field and transcranial conditions with a calibrated needle hydrophone (HNP0400, Onda, Sunnyvale, CA, USA). For accurate comparison between the trans-skull pressure between 0.8 and 1.5 MHz, transcranial focal pressure was used throughout the study. The transcranial focal pressure was obtained by locating the focus using hydrophone and 3D positioning system after placing a mouse skull harvested across age from 2-12 month in between the transducer and hydrophone. The vertical position of skull was carefully adjusted using pulse/echo so that the geometric focus of transducer was at approximately 4 mm inside the skull structure; lateral position of skull was also adjusted to a similar location used to target mouse hippocampus. At each frequency and conditions, a 30 cycle pulse was used, and the maximum peak-to-peak pressures were used to obtain calibration curve.

### Mathematical modeling of microbubble dynamics

To investigate the frequency-dependent behavior of microbubble (MB) dynamics in brain vasculature, we used a mathematical model that simulates ultrasound-driven MB oscillations, fluid transport across the vessel wall, and interstitial flow.^55^ MBs were modeled using a modified Rayleigh–Plesset equation that incorporates the non-linear effect of bubble oscillation due to its shell.^60,75–77^ To capture interactions with the vessel wall and surrounding fluid, the model was implemented in a finite element model comprising a lumen, vascular wall, and interstitial space, with the microbubble centrally positioned within the vessel. MB oscillations were coupled to the surrounding fluid via a pressure boundary condition, and vessel fluid dynamics was governed by the incompressible Navier–Stokes equations. The simulations were performed using the commercial finite element software, COMSOL (version 6.1, Burlington, MA, USA), where necessary equations were added using the Mathematics module.

### Microbubble preparation and characterization

The MBs used in this study were lipid-shelled MBs produced in-house (distearoylphosphatidylcholine (DSPC):1,2distearoyl-sn-glycero-3-phosphoethanolamine-N-[ma leimide(polyethylene glycol)-2000 (DSPE-PEG2000); 90:10 molecular ratio). To independently assess the impact of ultrasound frequency on the bioeffect and to ensure robust and consistent BBB opening, we utilized monodisperse MBs with a 5 μm diameter. These MBs were isolated through size-selective centrifugation of freshly activated polydisperse solution. For each 20 g mouse, a bolus injection of 50 μL containing 5 × 10^6^ MBs (equivalent to 2.5 × 10^8^ MBs/kg) was administered via the tail vein. To make sure the MB sizes and number of MBs selected were consistent using this method, the selected MB were counted and characterized by AccuSizer 770A Optical Particle Sizer (Santa Barbara, CA, USA) in triplicates before every MB-FUS treatment.

### Focused ultrasound treatment procedure

For the MB-FUS experiments, the animal was placed in the supine position with its head held by a stereotaxic frame designed in-house and attached to a 3D stage for fine positioning. To achieve precise FUS targeting, the hippocampal region of interest (ROI) in the right hemisphere was identified using ultrasound guidance for the 1.5 MHz setup and a needle-based alignment approach for the 0.8 MHz system. Target localization was guided by the skull contour and cross-referenced with anatomical landmarks from the Allen Mouse Brain Atlas and Allen Reference Atlas. To assess AAVs delivery, AAVs vectors were injected intravenously (i.v.) 1 minute before the MB-FUS treatment. The following exposure settings were employed: 1.5 MHz/0.8MHz therapy, 1 msec bursts, 5 Hz repetition rate, 320 kPa focal pressure; 5.2 MHz imaging, 1 cycle, 250 kPa, for a total treatment time of 2 minutes with concurrent i.v. tail vein administration of MB (2.5 × 10^8^ MBs/kg). Finally, the brains were harvested for further processing after performing transcardial perfusion with ice-cold PBS.

### Magnetic resonance imaging

To confirm the BBB opening, immediately after the sonication, the animals were injected with gadolinium contrast agent, Gd-HPDO3A (Prohance®, 0.5 μmol/g mouse body weight), and MRI was performed using a Bruker 11.7 Tesla small animal scanner (Bruker BioSpin MRI, Ettlingen, Germany) equipped with a cross coil configuration with a mouse body resonator for transmit and a mouse surface coil for receive. Images were acquired using ParaVision 360 (Bruker BioSpin MRI). Permeability of the BBB was determined with a T1 weighted (T1w) sequence (2D RARE sequence, RARE factor = 2, repetition time (TR) 250 ms, echo time (TE) 6.7 ms, 1 mm slice thickness, 1 mm interslice distance, 13 images, field of view (FOV) = 2 x 2 cm, matrix = 384 x 384, number of acquisitions (NA) = 6). Hemorrhage was assess using T2 weight MRI sequence (2D FLASH sequence, flip angle (FA) = 15°, repetition time (TR) 250 ms, echo time (TE) 15 ms, 1 mm slice thickness, 1 mm interslice distance, 13 images, field of view (FOV) = 2 x 2 cm2, matrix = 192 x 192, number of acquisitions (NA) = 4).

### Immunohistochemistry staining

Hematoxylin-eosin (H&E) staining was performed to examine tissue damage and safety. 20 µm thick frozen sections (Leica 3050 S Cryostat) were dehydrated beforehand and stained using a Leica Autostainer (ST5010). The sections were imaged with a 20x objective using a brightfield microscope (Eclipse Ti2, Nikon).

### AAV production

AAV9 vectors were produced by the UNC Vector Core (University of North Carolina). Briefly, AAVs were harvested 5 days after triple transfection in HEK293 cells by PEG precipitation of 3- and 5-days media and osmotic lysis of cell pellets. Crude AAVs were then purified by extraction from iodixanol density gradients and buffer exchanged into Dulbecco’s phosphate-buffered saline (DPBS). Viral titers were determined by qPCR on a woodchuck hepatitis virus post-transcriptional regulatory element (WPRE) present in all packaged AAV genomes.

### DNA extraction

To assess AAV9 vector biodistribution, mice were euthanized 21 hours post-treatment, and brains were extracted and bisected into sonicated and non-sonicated hemispheres on ice. The brain sample was immediately stabilized in Allprotect Tissue Reagent (Qiagen) until processing. DNA was isolated from tissue homogenates using the DNeasy Blood & Tissue Kit (Qiagen) according to the manufacturer’s protocol. DNA concentration and purity were determined by NanoDrop spectrophotometry (Thermo Fisher Scientific), and samples were diluted to a uniform concentration for qPCR analysis.

### RNA extraction and cDNA synthesis

To assess transgene expression, brain tissue was collected and immediately stabilized in RNA-protect tissue reagent (Qiagen) on ice. Total RNA was extracted from brain hemispheres using the RNeasy Midi Kit (Qiagen) following the manufacturer’s instructions. During this process, genomic DNA was removed by DNase. RNA integrity and concentration were assessed by NanoDrop spectrophotometry, and samples with an A260/A280 ratio between 1.8 and 2.0 were used for downstream analysis. Complementary DNA (cDNA) was synthesized from 500 ng-1 µg of total RNA using SuperScript IV VILO Master mix (ThermoFisher Scientific) according to the manufacturer’s protocol.

### Quantitative PCR (qPCR)

qPCR was performed using TaqMan-based assays with primers and probe designed to detect specific genome copies (tdTomato, PKM2, hBDNF, Syn1, Syp, and Dlg4) and normalized to the housekeeping gene, β-actin. Relative expression was calculated using the ΔΔCt method and expressed as fold-change relative to control groups. All reactions were performed in triplicate on a CFX96 (Bio-Rad) system using the following cycling conditions: 95°C for 10 min, followed by 40 cycles of 95°C for 15 s and 60°C for 1 min.

### Fluorescent reporter gene optical imaging

In vivo and ex vivo fluorescent imaging was performed using LAGO system (Spectral Instruments Imaging) with an exposure of 2s.

In vivo and ex vivo fluorescent imaging was performed using the LAGO imaging system (Spectral Instruments Imaging, Tucson, AZ) with excitation at 535 nm and emission at 589 nm (optimized for tdTomato fluorescence). Images were acquired with a 2-second exposure time, and fluorescence intensity was quantified using Aura, the manufacturer’s software. Regions of interest (ROIs) were drawn over the targeted hippocampal region, and fluorescence signal was quantified as maximum and mean radiance (photons/second/cm^2^/steradian) or total emission (photons/second).

### PET reporter gene imaging

Radiotracer synthesis was performed at the Stanford Cyclotron and Radiochemistry Facility (CRF). To noninvasively assess PKM2 expression, we employed [^18^F]DASA-10, a second-generation PET radiotracer with improved physicochemical and pharmacokinetic properties compared to [^18^F]DASA-23.^78^ [^18^F]DASA-10 was synthesized from nucleophilic displacement of the nitro group within 1-((2,3-dihydrobenzo[b][1,4]dioxin-6-yl)sulfonyl)-4-((2-fluoro-6-nitrophenyl)sulfonyl)piperazine, as previously described.^78^ The radiotracer ([^18^F]DASA-10; specific activity: 74.4 GBq/μmol) was administered via tail vein injection (∼5.55 MBq per mouse), followed by dynamic PET/CT imaging initiated at the time of injection and continued for 30 minutes.

Raw list mode data were reconstructed using 3D ordered-subset expectation maximization using maximum aposteriori (3D-OSEM/MAP) image reconstruction and converted to Standard Uptake Value (SUV). For dynamic image analysis of [^18^F]DASA-10, the 30-min list mode data was segmented into 20 static time frames (15 × 8, 60 × 8, 300 × 4; seconds x frames) and reconstructed as stated above. Quantitative PET image analysis was performed with Inveon Research Workplace (IRW) software after the co-registration of PET and CT images. Images were quantified by manually drawing ROI in the FUS treated area. PET images were quantified using standardized uptake values (SUV) to account for significant body weight variation across cohorts, as comparisons were made between healthy and 5xFAD mice ranging from 4 to 10 months of age. Images from 20-30 min post-injection were used for quantification to optimize signal-to-noise ratio. SUV max values represent the maximum voxel intensity within each ROI within this time inteval, not the temporal peak across the imaging window.

### Immunofluorescence staining and microscopy

For protein expression analysis, after the animals were euthanized at the time point based on their treatment protocols, the brains were harvested and were fixed with 4% paraformaldehyde overnight at 4°C. The next day, 20 µm sections were cut using a microtome (Leica 3050 S Cryostat).

To assess the biological effect induced by MB-FUS AAVs delivery, immunofluorescence staining was performed on the brain tissue. Tissues were prepared for staining by fixing in 4% paraformaldehyde at room temperature for 10 min (For sections requiring staining of intracellular markers (e.g., NeuN), they were permeabilized with 0.1% Triton X-100 in PBS for 5 minutes, subsequently). After washed with PBS, the sections were blocked for 1 hour at room temperature (2% Bovine Serum Albumin, 5% goat serum in PBS). It was then incubated with primary antibody diluted in 1% Bovine Serum Albumin (1:100) for 12 hours at 4°C. Next, the sections were incubated with secondary antibody diluted in 1% Bovine Serum Albumin (1:500) for 1 hour at room temperature. To stain the cell nucleus, samples were incubated with DAPI diluted in PBS (1:1000, 62248, Invitrogen) for 10 minutes after washing. Finally, the sections were rinsed with PBS to remove excess antibody, mounted with mounting medium (Prolong Glass Antifade Mountant, Lot# 2018752, Invitrogen), and covered with coverslips. Samples were cured with a mounting medium for 24 hours in dark at room temperature before imaging.

The sections were imaged with a 20x objective using a fluorescence confocal microscope (Leica DMi8 Inverted Microscope). The quantification of the fluorescence images was performed using ImageJ.

### Statistical analysis

All statistical analyses were performed using GraphPad Prism. P values P < 0.05 was considered statistically significant. (n.s. no significance, * P ≤ 0.05, ** P ≤ 0.01, ****P ≤ 0.0001). In the case of multiple comparisons, the p-values were adjusted using Bonferroni correction.

## Data availability

All data supporting the findings of this study are available within the article and its supplementary files. Any additional requests for information can be directed to, and will be fulfilled by, the corresponding authors. Source data are provided in this paper.

## Acknowledgments

We thank Dr. Frezghi Habte, Dr. Edwin Chang, and Laura Jean Pisani at the Stanford Center for Innovation in In Vivo Imaging (SCi3) for technical support with PET/CT, optical imaging, and MRI studies. [^18^F]DASA-10 production was carried out at the Stanford Cyclotron and Radiochemistry Facility (CRF). This study was supported by NIH grants R01 CA112356, R01 EB028646 and the Focused Ultrasound Foundation. Y.G. was partially supported by the Focused Ultrasound Foundation Lockhart Postdoctoral Fellowship.

## Author contributions

Y.G. and K.W.F. designed research; Y.G, J.F., and N.Z. performed FUS treatment; Y.G, J.A., N.Z., J.W.S., J.W., and B.L.J. performed PET/CT imaging, R.M. and C.B. developed and produced the PET radiotracer used in this study, Y.G., S.K.T and J.A. performed the bioassay and analyzed the data, B.L.J. and M.N.R. performed animal care, Y.G. and K.W.F. wrote the manuscript. All authors reviewed the manuscript and approved the final version.

