## Supplementary Information for "Ultrasound-Mediated Gene Therapy in Alzheimer’s Disease Validated through In Vivo PET Imaging"

### Supplementary Methods

#### 1. Optimization of microbubble size for AAV delivery to the brain with MB-FUS.

To study the effect on microbubble size on BBB opening with MB-FUS, we conducted MB-FUS experiments using same number of monodispersed lipid shell MBs that were separated into different size, 2 µm and 5 µm ($2.5\times10^{8}$ MB/kg) (Suppl. Fig. 1 A-B). In this experiment, to mimic the AAV delivery, we assessed the delivery of 70 kDa fluorescently labeled dextran particles (14 nm) in the brain. The MRI contrast enhanced imaging and ex vivo fluorescent imaging shows significantly higher enhancement for 5 µm MBs as compared to 2 µm MBs, suggesting that the larger MB has a stronger influence in changing vessel permeability (Suppl. Fig. 1 C-E).

#### 2. Optimization of ultrasound pressure for for AAV delivery to the brain with MB-FUS.

To determine the most effective ultrasound pressure within the safety range, we tested focal pressures accounting for insertion loss ranging from 350 kPa to 550 kPa peak negative pressure with 5µm microbubble. The acoustic emission showed stronger harmonic level as pressure increases (Suppl. Fig. 2 B-D). However, high level of broadband emissions (MB collapse) were observed during the sonication for pressure higher than 420kPa, which leads to hemorrhage in the brain (Suppl. Fig. 2 D-G).

### Supplementary Figures

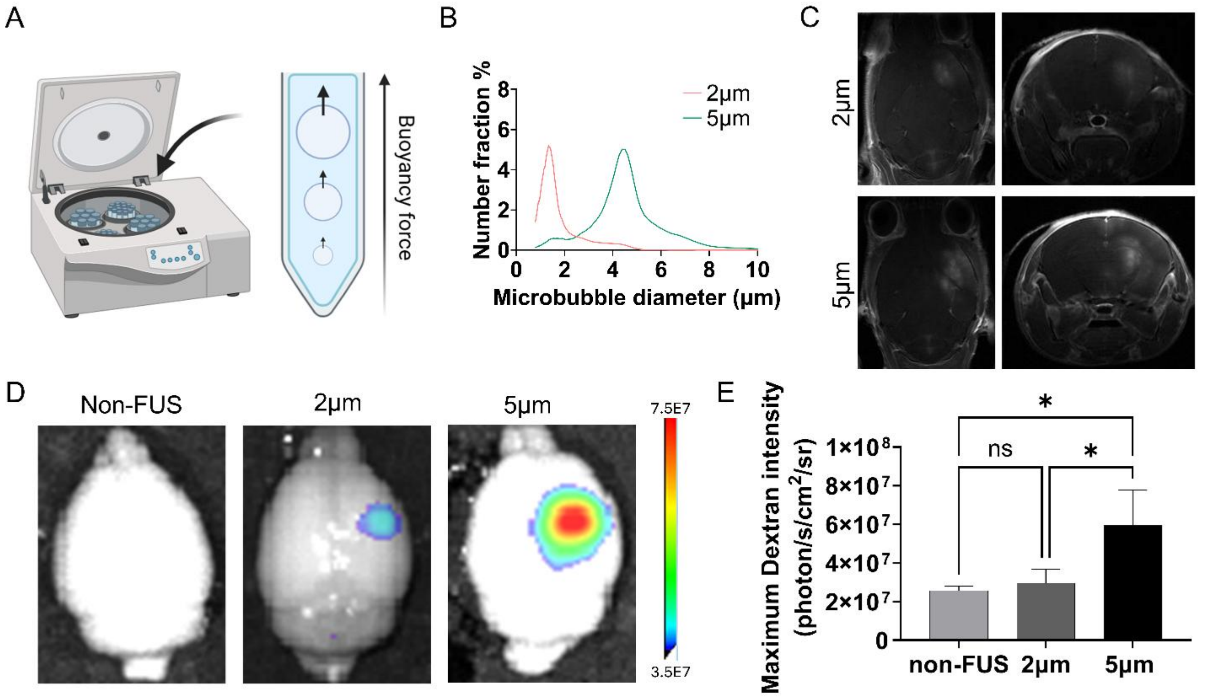


**Suppl. Fig. 1.** A) schematics of microbubble size isolation method. B) Size distribution for 2µm and 5µm microbubble. C) Representative contrast enhanced T1-weighted MR image after MB-FUS treatment with 2µm and 5µm microbubble. D) Representative ex vivo images of dextran particle delivery in the brain. E) Quantification of ex vivo images of dextran particle delivery in the brain (n=3). P values were determined by one-way analysis of variance (ANOVA). Plots show means ±S.D. n.s. no significance, * $P \leq 0.05$,

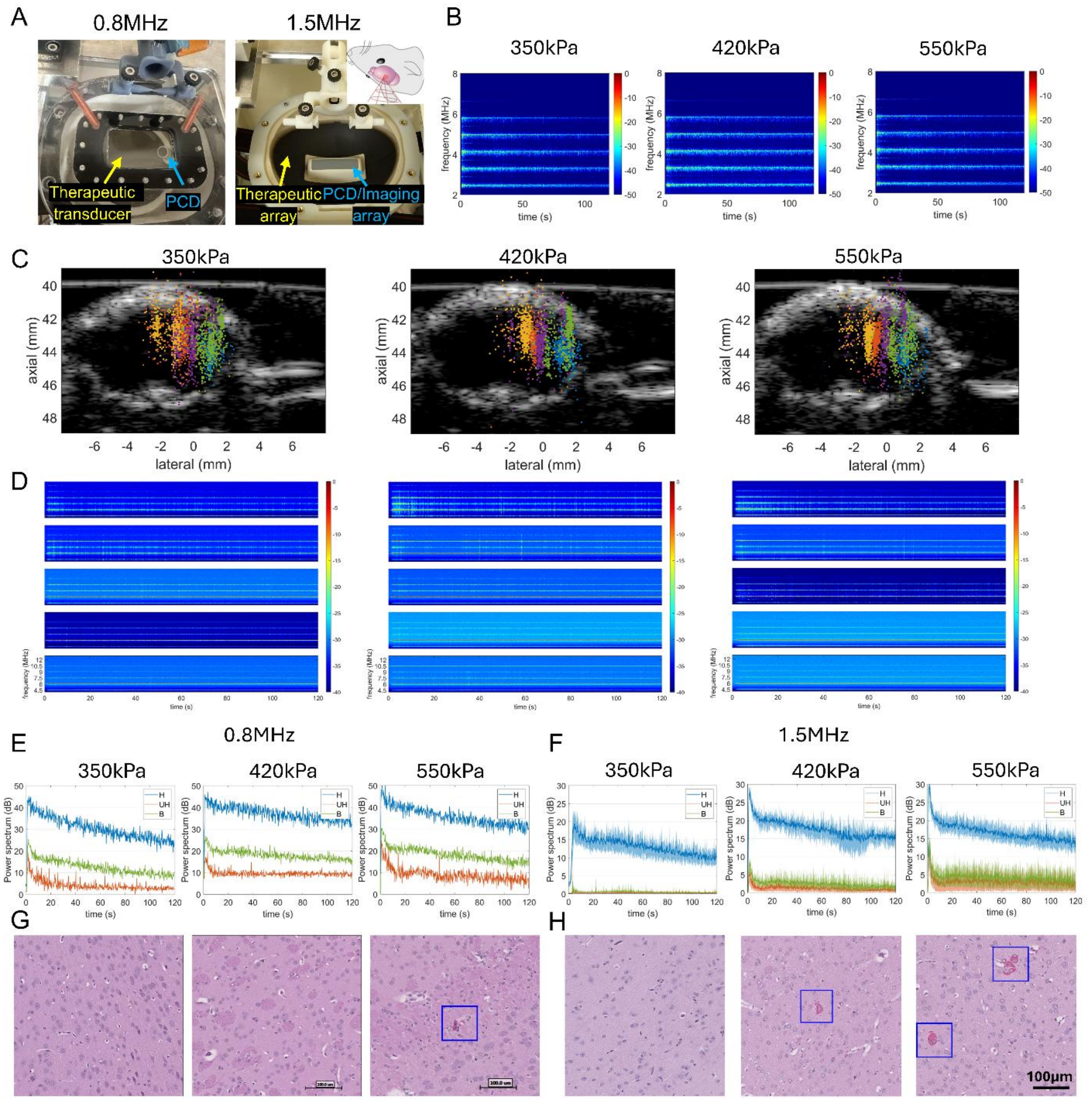


**Suppl. Fig. 2.** A) 0.8MHz (left) and 1.5MHz (right) focused ultrasound setup. B) Spectrogram during sonication using 0.8 MHz frequency at 350 kPa, 420 kPa and 550 kPa pressure. C) Post-FUS passive acoustic map localizing acoustic emissions from MBs for the entire treatment using 1.5 MHz frequency at 350 kPa, 420 kPa and 550 kPa pressure. D) Spectrogram during sonication using 1.5 MHz frequency at 350 kPa, 420 kPa and 550 kPa pressure. E) Power levels of acoustic emissions during treatment using 0.8 MHz frequency at 350, 420, and 550kPa peak negative pressure. F) Power levels of acoustic emissions during treatment using 1.5 MHz frequency at 350, 420, and 550kPa peak negative pressure. G) Representative H&E staining images after MB-FUS treatment using 0.8 MHz frequency at 350, 420, and 550 kPa peak negative pressure. H) Representative H&E staining images after MB-FUS treatment using 1.5 MHz frequency at 350, 420, and 550 kPa peak negative pressure. Abbreviations: H: harmonics, UH: ultraharmonics, B: broadband.

29

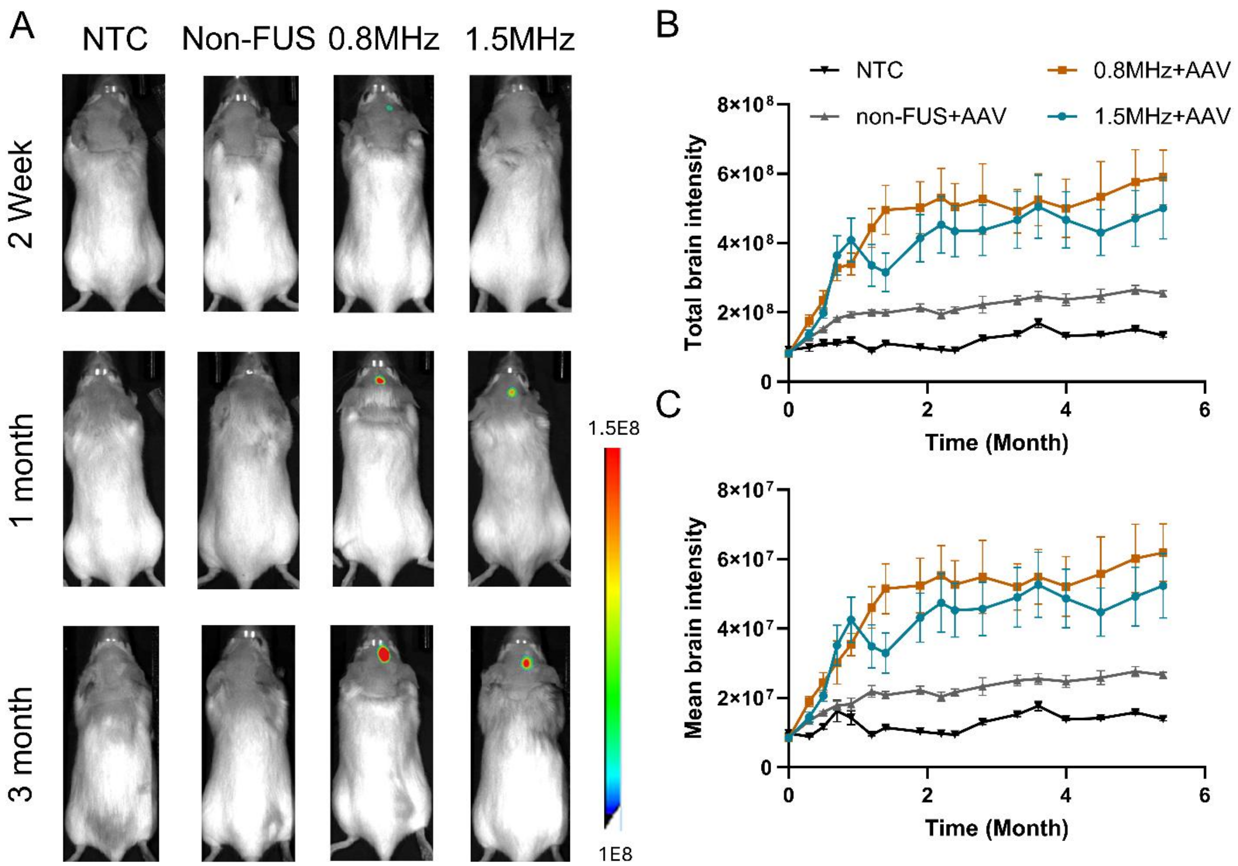


**Suppl. Fig. 3.** A) Representative whole-body supine imaging showing AAV9-tdTomato expression in the brain at 2 weeks, 1 month, and 3 months post-injection. B) Quantification of in vivo imaging of total tdTomato expression in the brain over a 6-month period following treatment. C) Quantification of in vivo imaging of mean tdTomato expression in the brain over a 6-month period following treatment.

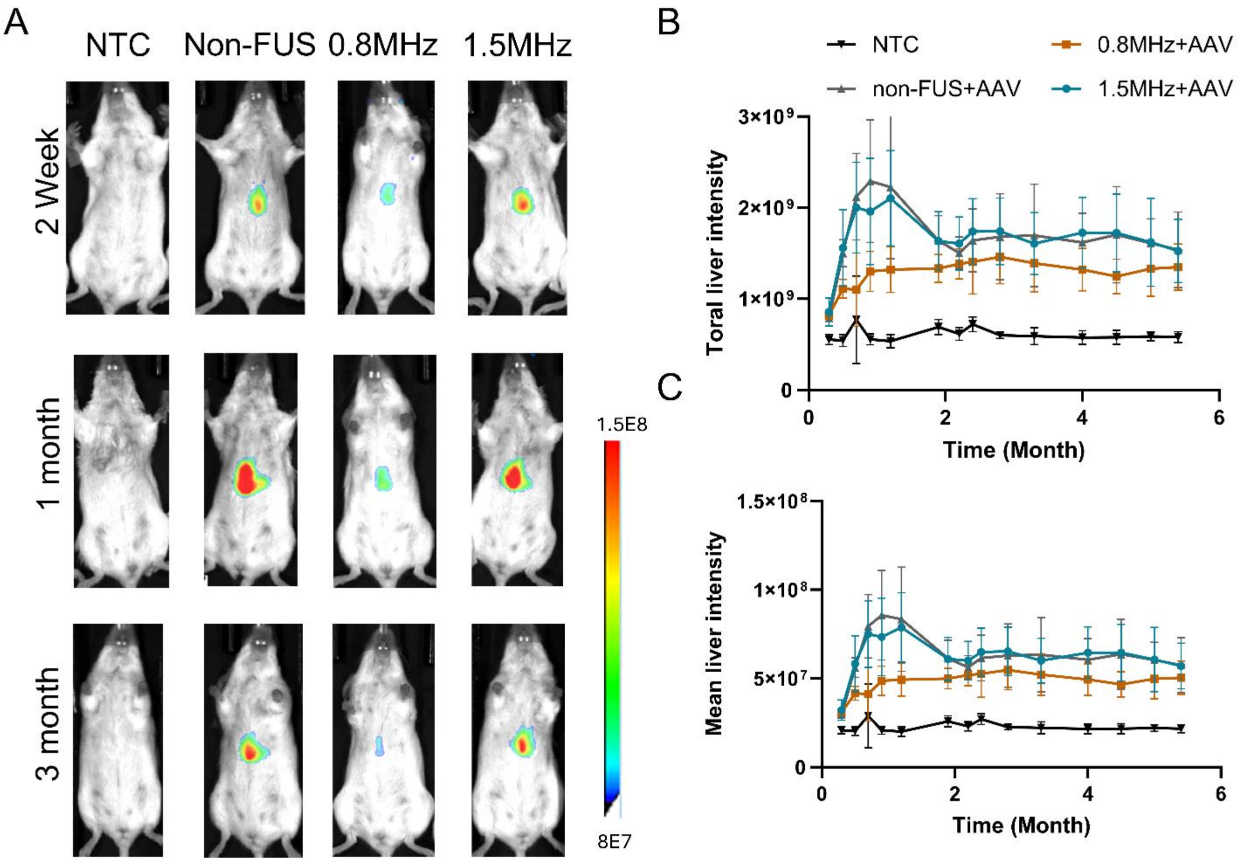


**Suppl. Fig. 4.** A) Representative whole-body prone imaging showing AAV9-tdTomato expression in the liver at 2 weeks, 1 month, and 3 months post-injection. B) Quantification of in vivo imaging of total tdTomato expression in the liver over a 6-month period following treatment. C) Quantification of in vivo imaging of mean tdTomato expression in the liver over a 6-month period following treatment.

31

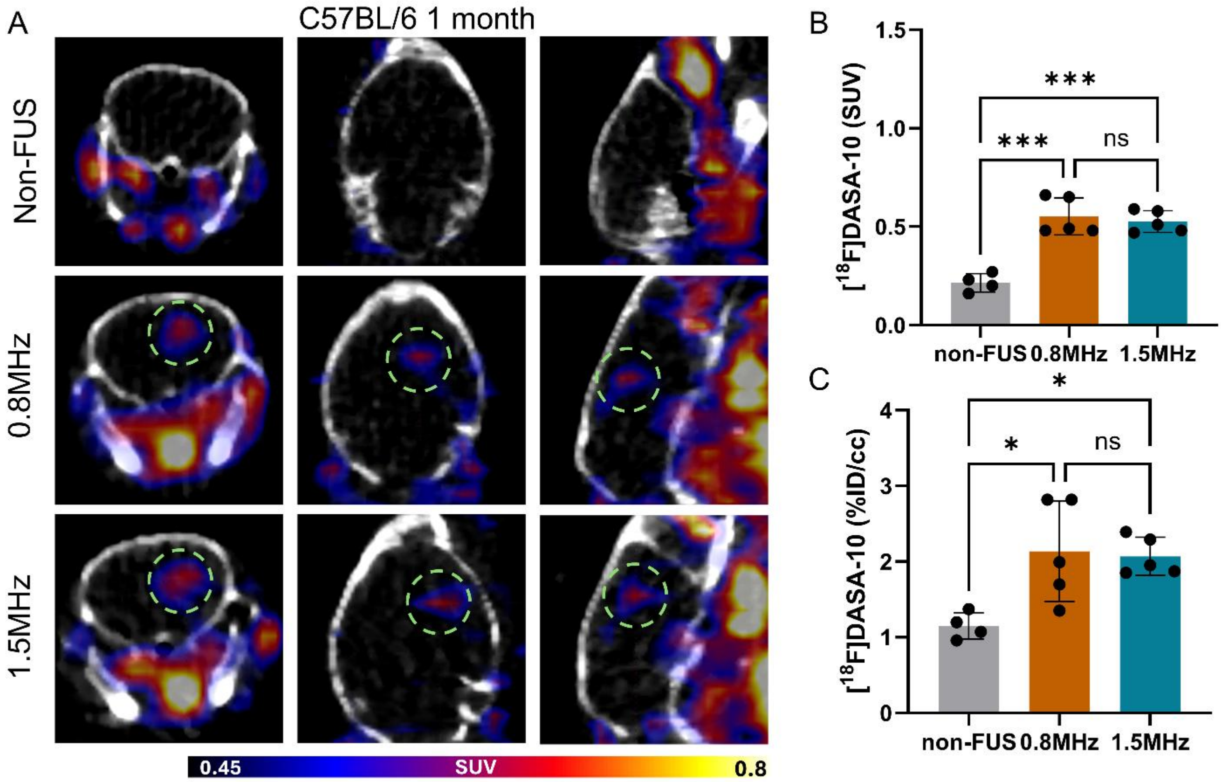


**Suppl. Fig. 5.** A) Representative PET/CT images (coronal, axial, sagittal) obtained immediately following [$^{18}$F]DASA-10 administration, 1 months after MB-FUS-assisted AAV9-PKM2 delivery in healthy mice. Regions of MB-FUS treated region are marked with dotted green circles. B) Image-based quantification of standardized uptake values (SUVs) in regions of interest across treatment groups in healthy mice following [$^{18}$F]DASA-10 injection. C) Quantification of [$^{18}$F]DASA-10 uptake (%ID/cc) in regions of interest 1 month following AAV administration in healthy mice. P-values were determined by one-way analysis of variance (ANOVA) and were adjusted using Bonferroni correction. Plots show mean ± S.D. (n = 4-5) n.s. no significance P >0.05, *P ≤ 0.05, *** P ≤ 0.001.

32

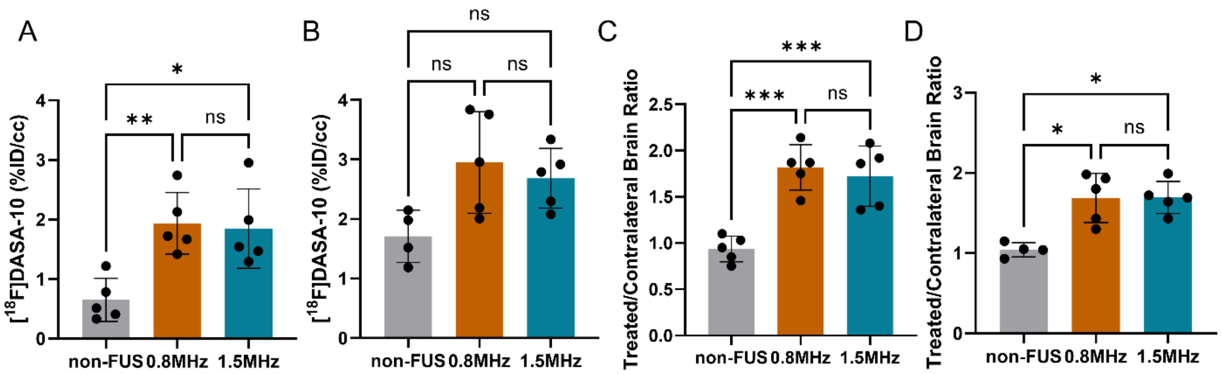


**Suppl. Fig. 6.** A) Quantification of [$^{18}$F]DASA-10 uptake (%ID/cc) in regions of interest 7 month following AAV administration in healthy mice. B) Quantification of [$^{18}$F]DASA-10 uptake (%ID/cc) in regions of interest 7 month following AAV administration in 5xFAD mice. C) Quantification of DASA-10 uptake ratio between FUS-treated and contralateral hippocampus at 7 months following AAV administration in healthy mice. D) Quantification of DASA-10 uptake ratio between FUS-treated and contralateral hippocampus at 7 months following AAV administration in 5xFAD mice. Note that contralateral uptake may reflect both background tracer signal and PKM2 expression from transduced cells due to interhemispheric AAV transport following FUS treatment. P-values were determined by one-way analysis of variance (ANOVA) and were adjusted using Bonferroni correction. Plots show mean ± S.D. (n = 4-5) n.s. no significance, $P > 0.05$, $^*P \leq 0.05$, $^{**}P \leq 0.01$, $^{***}P \leq 0.001$.

33

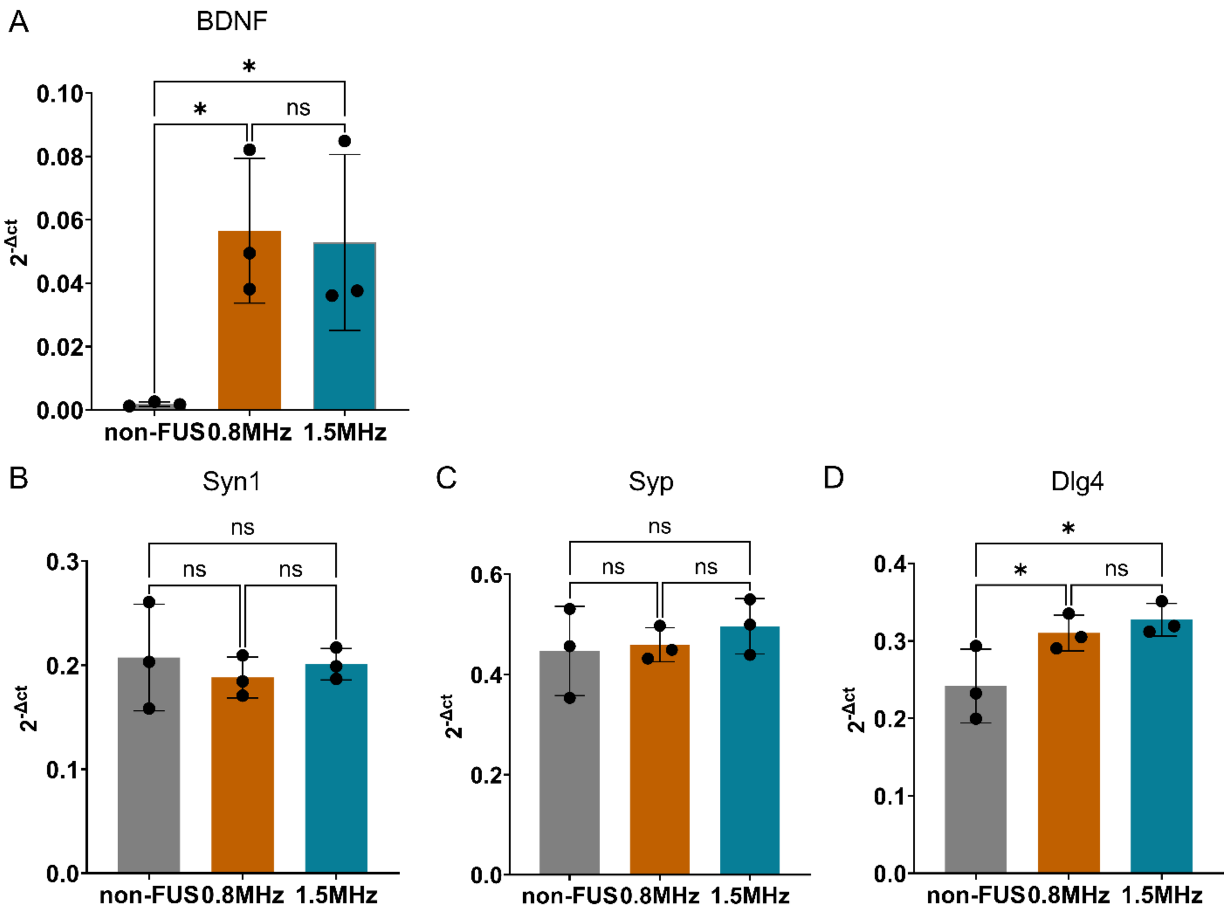


**Suppl. Fig. 7.** A) RT-qPCR quantification of BDNF mRNA levels in the treated hemisphere of C57BL/6 mice 1 month following MB-FUS-mediated AAV9-BDNF delivery. RT-qPCR quantification of the B) presynaptic marker synapsin 1 (Syn1), C-D) postsynaptic markers synaptophysin (Syp) and PSD-95 (Dlg4) one month post treatment. P-values were determined by one-way analysis of variance (ANOVA) and were adjusted using Bonferroni correction. Plots show mean ± S.D. (n = 3) n.s. no significance $P > 0.05$, $^*P \leq 0.05$.
